# Evolution of transient RNA structure-RNA polymerase interactions in respiratory RNA virus genomes

**DOI:** 10.1101/2023.05.25.542331

**Authors:** Charlotte Rigby, Kimberly Sabsay, Karishma Bisht, Dirk Eggink, Hamid Jalal, Aartjan J.W. te Velthuis

## Abstract

RNA viruses are important human pathogens that cause seasonal epidemics and occasional pandemics. Examples are influenza A viruses (IAV) and coronaviruses (CoV). When emerging IAV and CoV spill over to humans, they adapt to evade immune responses and optimize their replication and spread in human cells. In IAV, adaptation occurs in all viral proteins, including the viral ribonucleoprotein (RNP) complex. RNPs consists of a copy of the viral RNA polymerase, a double-helical coil of nucleoprotein, and one of the eight segments of the IAV RNA genome. The RNA segments and their transcripts are partially structured to coordinate the packaging of the viral genome and modulate viral mRNA translation. In addition, RNA structures can affect the efficiency of viral RNA synthesis and the activation of host innate immune response. Here, we investigated if RNA structures that modulate IAV replication processivity, so called template loops (t-loops), vary during the adaptation of pandemic and emerging IAV to humans. Using cell culture-based replication assays and *in silico* sequence analyses, we find that the sensitivity of the IAV H3N2 RNA polymerase to t-loops increased between isolates from 1968 and 2017, whereas the total free energy of t-loops in the IAV H3N2 genome was reduced. This reduction is particularly prominent in the PB1 gene. In H1N1 IAV, we find two separate reductions in t-loop free energy, one following the 1918 pandemic and one following the 2009 pandemic. No destabilization of t-loops is observed in the IBV genome, whereas analysis of SARS-CoV-2 isolates reveals destabilization of viral RNA structures. Overall, we propose that a loss of free energy in the RNA genome of emerging respiratory RNA viruses may contribute to the adaption of these viruses to the human population.

## Introduction

RNA viruses, such as influenza A viruses (IAV) or coronaviruses (CoV), are important pathogens that cause mild to severe respiratory disease in humans. RNA virus spread among humans occurs in seasonal epidemics, outbreaks of emerging viruses, or occasional pandemics [1]. Seasonal epidemics are caused by viruses that are largely adapted to humans and to which humans have pre-existing immunity, such as the seasonal IAV strains H3N2 and H1N1 or CoV strains NL63 and 229E. In contrast, outbreaks and pandemics are caused by emerging RNA viruses to which humans lack pre-existing immunity [2]. Emerging viruses spill over from an animal reservoir to humans, and typically have poor human-to-human transmission. Examples of these are the IAV H5N1 and H7N9 strains. Pandemic viruses can efficiently spread from human-to-human, due to adaptations that they acquired via mutation and natural selection, or through reassortment or recombination with a human-adapted virus. In recent history, IAV subtypes H1N1, H2N2 and H3N2 have caused pandemics in 1918, 1957, 1968 and 2009, while the CoV severe acute respiratory CoV 2 (SARS-CoV-2) caused the COVID-19 pandemic [3].

Analysis of IAV strains has shown that human-adaptive mutations occur in nearly all viral proteins, but in particular in the IAV surface glycoproteins hemagglutinin (HA) and neuraminidase (NA); the proteins involved in viral replication polymerase basic 1 and 2 (PB1 and PB2), polymerase acidic (PA) and the nucleoprotein (NP); and the protein involved in modulating host-immune responses, non-structural 1 (NS1) [4–16]. During the SARS-CoV-2 pandemic, adaptive mutations have been observed primarily in the spike (S) glycoprotein, and potentially important mutations have been reported for the nucleocapsid protein, envelope protein, and non-structural proteins 2, 3, 4, 10, 12 and 13 [17–23]. Research indicates that mutation in the glycoproteins improve host receptor binding, virus transmission, pH stability and host immune evasion, whereas mutations in the viral replication complex improve viral RNA synthesis [17,24–31]. In addition to mutations in the viral proteins, nucleotide changes and deletions have been observed in IAV and SARS-CoV-2 genome RNA structures located in untranslated regions (UTRs), indicating that changes at the RNA structure level may also play a role in the adaptation of RNA viruses [32,33].

Secondary RNA structures in RNA virus genomes play critical roles in IAV and CoV infection cycles and perform functions in packaging, translation, and viral replication. In addition, RNA structures can be the target of therapeutics [34]. Secondary RNA structures can form between neighboring sequences, as well as involve long-range interactions, such as duplex formation between the 5ʹ and 3ʹ UTRs of various viruses [35,36]. Disruption of some RNA structures affects RNA virus growth, including packaging signals, however several IAV RNA structures have a large degree of variability among strains [37–39]. Presently, our understanding of the role of RNA secondary structures as well as mutations in these structures during RNA virus replication and adaptation is limited, but it seems likely that RNA structures that negatively impact viral replication in a new host may be modified or removed in time.

In this study, we aimed to gain insight into the evolution of template loops (t-loop), a transient RNA secondary structure that folds effectively “around” the viral RNA polymerase and affects IAV replication efficiency on short viral RNA templates [40]. In the t-loop model, we envision the RNA polymerase activity as an object on a tablecloth: when the enzyme synthesizes RNA, each translocation of the RNA polymerase changes the local RNA template “landscape”. Some structures may be (partially) perturbed and others RNA may form as their folding energy now becomes favorable. We can use a sliding-window algorithm to calculate the deltaG for RNA structures upstream, downstream and around the RNA polymerase along an RNA sequence. Currently, the effect of t-loops in full-length viral genome (segments) cannot be studied due to the lack of appropriate biochemical assays. However, we hypothesized that if strong t-loops are present in the viral genome and they affect the efficiency of viral RNA synthesis, they would be destabilized over time to improve viral fitness. These destabilizations would appear as synonymous nucleotide changes in primary sequence data. We expected that these putative t-loop destabilizations would occur in emerging IAV strains, because they would benefit most from improvements in viral replication. In addition, we hypothesized that after t-loop destabilization, continued adaption of the viral replication complex in a t-loop-less genome would make the viral RNA polymerase sensitive to the re-introduction of t-loops. Hence, we expected human-adapted IAV to be more sensitive to t-loops structures than recently emerged/less human-adapted viruses, and that reassortment events between human and avian-adapted IAV would impose strong selections on strong t-loops, which would be apparent from sequence data.

To test the above hypotheses, we performed cell culture assays as well as analyses of IAV, IBV and SARS-CoV-2 sequence data obtained through surveillance. We observe that the sensitivity of H3N2 IAV RNA polymerases to t-loops increased between 1968 and 2017, but that the total free energy (ΔG) of t-loops in the IAV H3N2 genome was linearly increased, and thus the overall stability reduced. Reductions in t-loop stability are also observed in H1N1 IAV following the 1918 and 2009 pandemics. No destabilization of t-loops is observed in the IBV genome, which has been circulating in humans for decades, but an analysis of SARS-CoV-2 isolates reveals destabilization of t-loops as well. Overall, we propose that the ΔG of RNA structures in the RNA genome of emerging respiratory RNA viruses may play a role in the adaption of these viruses to humans.

## Results

### T-loops differentially affect IAV H3N2 RNA polymerase activity and IFN induction

We previously identified t-loops as RNA elements that can reduce the replication efficiency of H1N1 and H5N1 IAV RNA polymerases on 71-nt long RNA templates called mini viral RNAs (mvRNAs) [40]. In our previous observations, mvRNAs containing a t-loop in the first half of the template were more likely to trigger IFN promoter activity than mvRNAs without a t-loop in the first half of the template [40]. So far, the sensitivity of the RNA polymerase of emerging and adapted IAV viruses to t-loops has not been directly compared. The IAV H3N2 genome has been continuously circulating in humans for 55 years, making the replication complexes of H3N2 IAVs ideal for performing a direct comparison between an emerged and adapted IAV strain. To perform the comparison, we transfected plasmids expressing the PB1, PB2, PA and NP proteins of the A/WSN/1933 (abbreviated as WSN), A/Netherlands/1968 (abbreviated as H3N2-1968), and A/Netherlands/2017 (abbreviated as H3N2-2017) RNA polymerases into HEK 293T cells along with an IFN-promoter driven luciferase reporter plasmid and a TK-promoter-driven *Renilla* luciferase control plasmid. In addition, we provided a cellular RNA polymerase I-driven plasmid expressing a 71-nt long template without a t-loop in the first half of the template (NP71.1) and a 71-nt long template containing a t-loop in the first half of the template (NP71.11). These templates were extensively tested previously and used to show that the IFN promoter activity is indicative of poor RNA polymerase activity [40]. Western blot analysis showed efficient expression of the viral proteins (Fig. 1) Measurement of the IFN promoter activity showed no difference in the ability of the WSN and H3N2-1968 RNA polymerase to induce IFN promoter activity. However, the H3N2-2017 RNA polymerase induced an almost 8-fold higher IFN promoter activity (Fig. 1), suggesting that it had a higher sensitivity to the t-loop structure in the template than the WSN and H3N2-1968 RNA polymerase.

**Figure 1.**
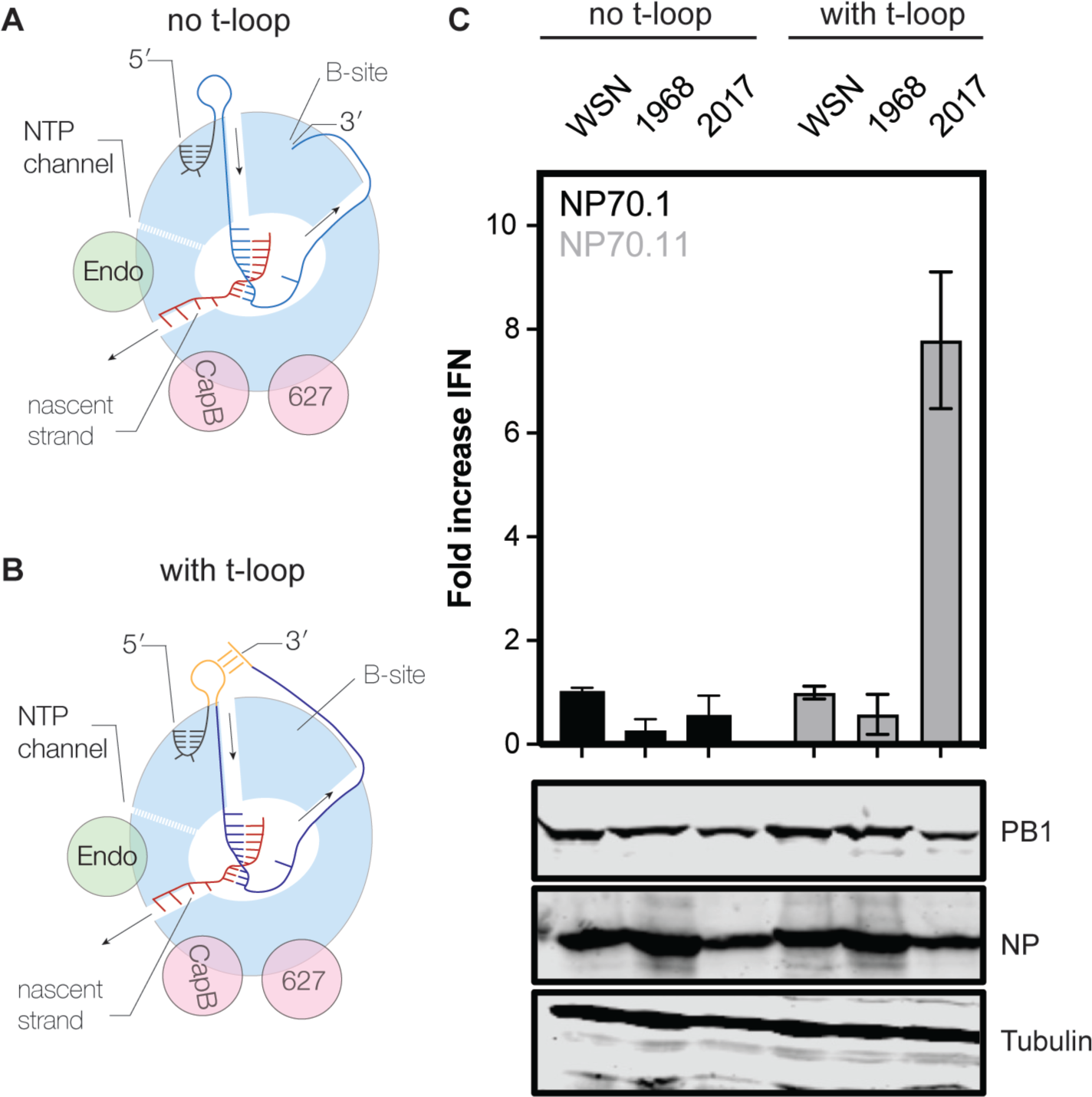
Interferon beta promoter activity in influenza A virus H3N2 RNP reconstitution assay. **A**) Model of the elongation complex of the IAV RNA polymerase, in which the 3ʹ is bound by the B-site on the side of the RNA polymerase. Arrows indicate movement of template and product RNA through the RNA polymerase. **B**) Model of the elongation complex of the IAV RNA polymerase, in which the 3ʹ base-pairs with a sequence located upstream of the RNA polymerase. The formation of such a loop in the template leads to reduced RNA polymerase activity and innate immune activation when the loop forms in the first half of the template. **C**) Fold change in *IFNB* promoter activation following the replication and transcription of an mvRNA containing no t-loop in the first half of the template (NP71.1) and an mvRNA containing a t-loop in the first half of the template (NP71.11). Fold change in *IFNB* promoter activation was normalized to the IAV WSN H1N1 control for each template. Bottom panels show western blot analysis of NP and PB1 expression for the analyzed IAV H3N2 replication complexes. Tubulin was used as loading control.

### Initial analysis of IAV H3N2 t-loop stability

The above result suggested that the RNA polymerase of a recent IAV H3N2 isolate is more sensitive to t-loops than an IAV H3N2 isolate from 1968. We hypothesized that any t-loops in the 1968 IAV H3N2 genome would be absent in the 2017 IAV H3N2 genome if they negatively affected viral replication. To explore this, we focussed on an initial set of genome sequences of IAV H3N2 isolates from 1968, 1972, 1982, 1993, 2003, 2008, 2014, and 2017. These isolates capture the full IAV H3N2 tree (Fig. 2A) and did not show a particular trend in the variation of the GC content per genome (Fig. 2B).

**Figure 2.**
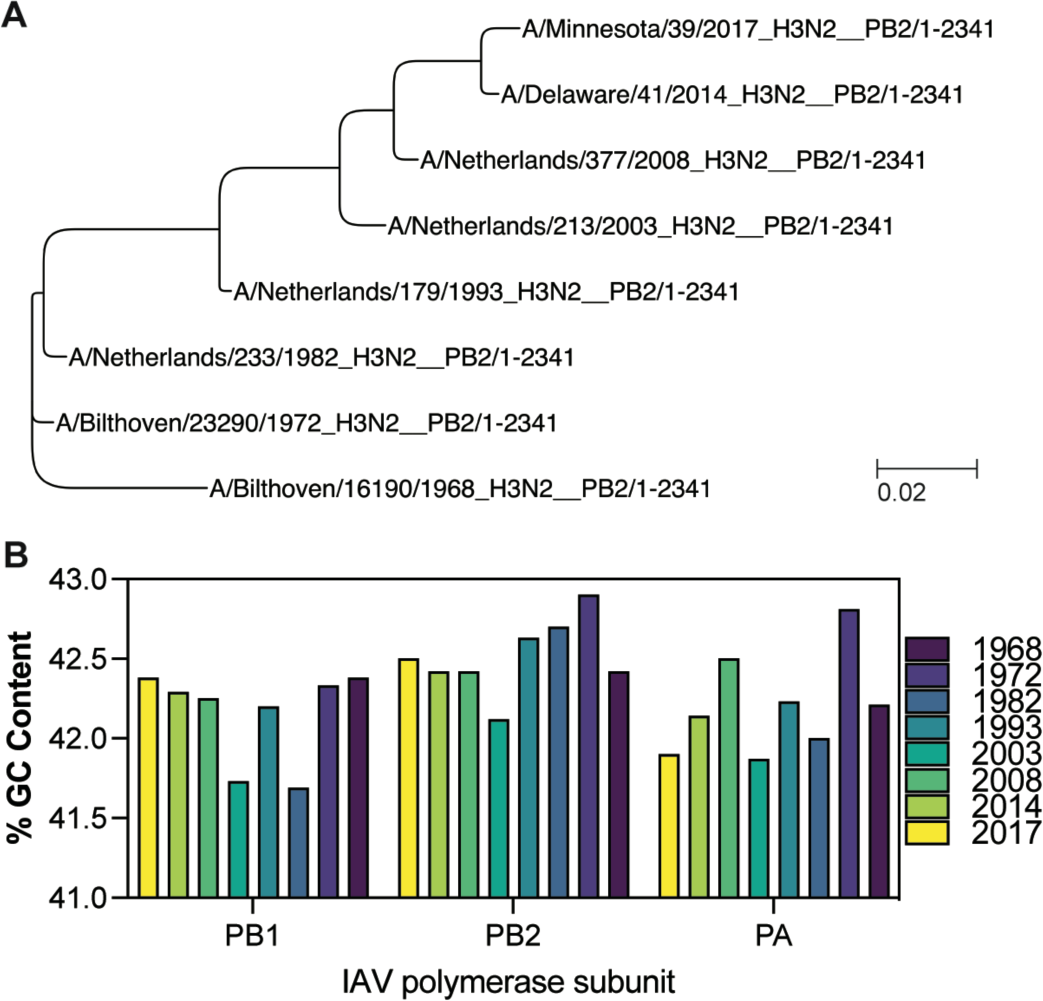
Phylogeny and GC content of IAV H3N2 genomes between 1968 and 2017. A) Phylogenetic analysis of IAV H3N2 based on the PB2 nucleotide sequences of isolates from 1968, 1972, 1982, 1993, 2003, 2008, 2014 and 2017. Bar represents degree of nucleotide change. B) GC content (%) for the genomes shown in Fig. 2A.

Next, we analyzed the putative RNA structure content in the IAV H3N2 genome. To this end, we modified the t-loop analysis that we described previously [40] to account for NP binding up and downstream of the IAV RNA polymerase during replication and RNP assembly (Fig. 3A and B). Specifically, we used a sliding window analysis, in which a sequence covered by the 20-nt footprint of the IAV RNA polymerase is blocked from participating in secondary RNA structure formation [41,42]. We also assumed that NP has a footprint of 24 nt, and that the separation of the template RNA from the downstream or upstream NP in the RNP makes 24 nt available for base pairing (Fig. 3B) [43]. With both an upstream and downstream NP removed, t-loop as well as any other secondary RNA structure formation can be calculated between the two 24-nt of single-stranded RNA (Fig. 3B). The sliding window was started at the 3’ end of the viral genome and moved in 1-nt intervals along the length of the template RNA.

**Figure 3.**
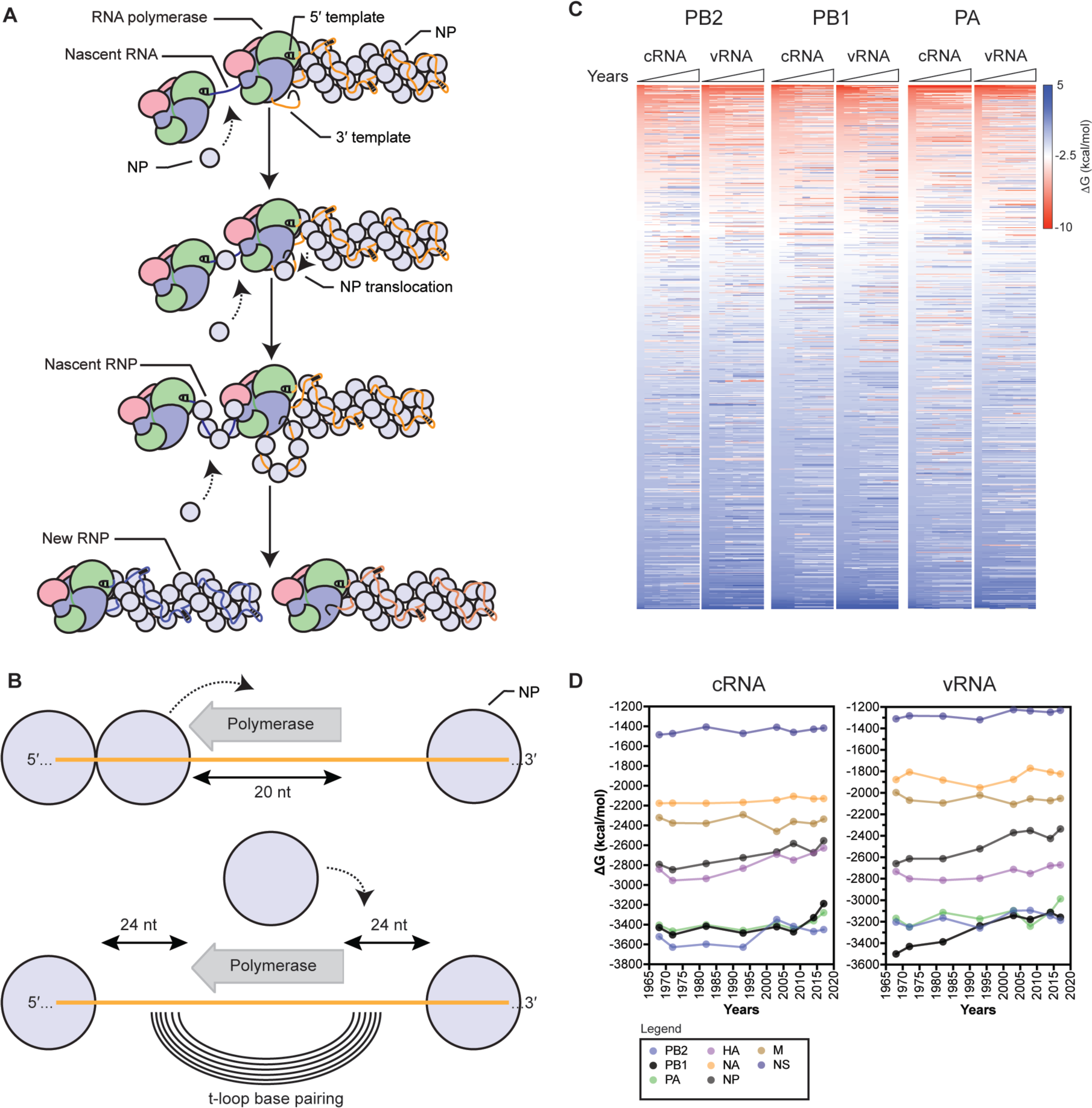
Model for RNP assembly during replication and analysis of influenza A virus H3N2 genomes. A) Model of IAV replication and RNP assembly following the recruitment of a second RNA polymerase and newly synthesized NP molecules. B) Model of t-loop formation in an IAV RNP that is the basis of this manuscript. The template is shown in orange, with the footprint of the RNA polymerase indicated as a grey arrow. Base pairing between the up- and downstream sequences is illustrated as concentric lines. The footprint of the NP molecules, 24 nt, is also indicated. C) Analysis of the t-loop ΔG in the IAV H3N2 genome segments 1-3. Analysis was performed on the IAV H3N2 genomes of Fig. 2 and the ΔG sorted from negative to positive on the 1968 H3N2 values. See Table S1 for unsorted data. Scale bar indicates t-loop ΔG in kcal/mol. D) Total t-loop ΔG in IAV genome segments for the IAV H3N2 genomes of Fig. 2. See Table S1 for data.

Before discussing our findings, we note a few caveats to our approach. Research has shown that the placement of NP along the viral genome is heterogeneous, and that various secondary RNA structures are present in IAV genome segments *in virio* [44,45]. NP dissociation downstream or upstream of an existing RNA structure may make additional nucleotides available for t-loop formation. However, the stability of such a secondary RNA structure is likely such that it will not spontaneously unwind and contribute to t-loop formation. In addition, if spontaneous unwinding were to occur, any nucleotides placed further away from the RNA polymerase template channels are unlikely to form a tight t-loop around the RNA polymerase and will thus unlikely contribute to the modulation of RNA polymerase processivity. For simplicity, we have excluded the above two possibilities from our analysis, but point them out because we appreciate that the (transient) RNA structure landscape is likely more complex than we can currently model.

Analysis of the first three IAV H3N2 genome segments showed a location-specific loss of, as well as an increase in t-loop stability across the segments in both the cRNA and vRNA sense (Fig. 3C; Table S1). We also noticed that positive ΔG values were present and increased in some locations. Together, these observations suggest that the stability of t-loops in the IAV H3N2 genome varies over time and per location. To get a better sense of the impact of changes in t-loop stability across each segment in time, we calculated the total t-loop ΔG per segment, per IAV H3N2 isolate. To avoid introducing biases (e.g., varying number of data points, excluding parts of some segments for some isolates, etc), we included both the positive and negative ΔG values in this analysis. Interestingly, we observed that of the three RNA polymerase encodings segments, in particular the PB1 vRNA segment showed a trend corresponding with t-loop destabilization (Fig. 3D), while other segments showed only a modest variation in time. This trend in PB1 was also present if we analyzed the negative t-loop ΔG values only (Fig. S1).

To investigate whether the observed trends were generated by chance, we simulated mutations in the IAV H3N2 isolate from 1968, with frequencies of 3.2 x 10^-3^ and 2.5 x 10^-4^ per nucleotide per year as the upper and lower bounds of the whole genome mutation rate [46,47]. We performed the mutation simulations for the PB1 and PB2 segments for 3 consecutive decades, while maintaining the codon sequence. Next, we calculated the t-loop ΔG for the *in silico* variants in both the cRNA as vRNA sense. As shown in Fig. S2, the t-loop ΔG of the PB1 segment deviated from the *in silico* variants, trending towards less negative ΔG values, whereas the PB2 t-loop ΔG values overlapped with the *in silico* variants. However, the observed patterns were not significantly different for this small sample size.

### The IAV H3N2 shows a progressive loss of t-loop stability in PB1 segment

We next extended our dataset using 50 full-length IAV H3N2 sequences for each of the above years (Table S2). We first analyzed the total minimum free energy (MFE; which attempts to describe large RNA structures and structure forming up and downstream of the RNA polymerase) in segment 1 (PB2), segment 2 (PB1) and segment 3 (PA) as well as the total t-loop stability per IAV genome segment. In line with our initial analysis above, we observed clear segment-specific trends (Fig. 4). The total MFE followed a progressive increase in MFE stability (decrease in ΔG) for the PB2 segment, whereas the PB1 and PA segments demonstrated a loss of MFE stability (Fig. 4). In contrast, the t-loop stability was reduced for all three RNA polymerase segments across the 55 years analyzed. Compared to the *in silico* variants (Fig. S2), the PB1 change in ΔG was significantly different (p<0.001) The steepest increase in t-loop ΔG was observed for the PB1 segment between 1968 and 2008. An apparent plateau in PB2 and PB1 t-loop stability followed 2008, but additional (future) years would need to be analyzed to confirm the start of a new trend. For simplicity, we fit the trends with a linear regression (p <0.0001). In general, both the vRNA and cRNA sense showed similar directions in their increase or decrease in MFE or t-loop ΔG. However, the direction in MFE and t-loop ΔG change was not consistent, as was particularly evident in the PB2 segment (Fig. 4).

**Figure 4.**
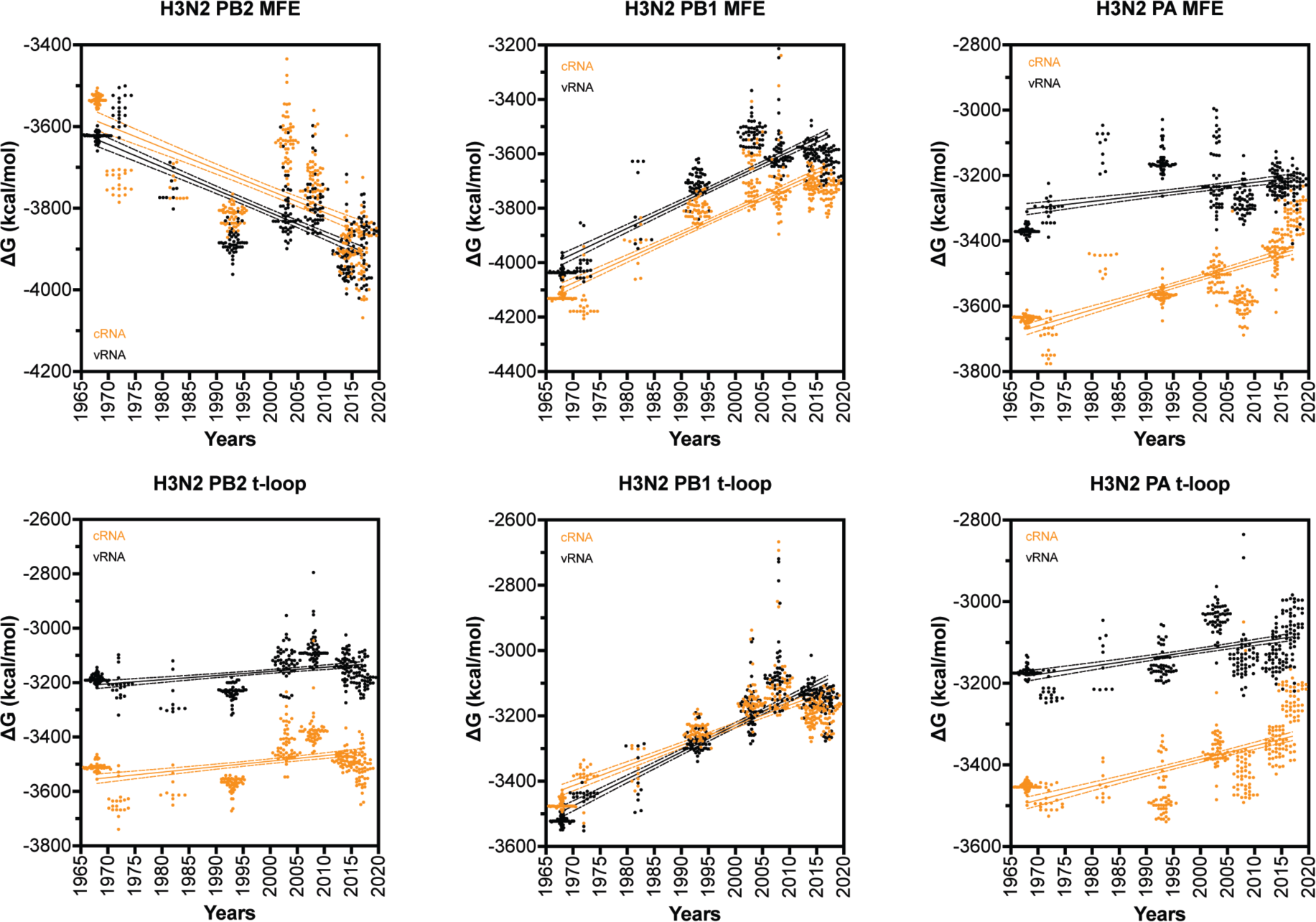
Total MFE and t-loop free energy analysis for IAV H3N2 isolates between 1968 and 2017. Analysis of the predicted MFE and t-loop content of the cRNA (orange) and vRNA (black) is shown. For each year, the first 50 full-length, unique sequences identified (where available) were analyzed and plotted. Data points were fit with linear regression. Dotted line indicates 95% confidence interval.

### Changes in t-loop stability occur independently of codon mutation

Other RNA elements (e.g., packaging signals) may contribute or drive the changes in t-loop stability observed in Fig. 3 and 4, but presently it is unclear where they are located and how much each putative RNA element contributes to the IAV infection cycle. However, it is well established that IAVs proteins adapt through amino acid mutation, including the polymerase subunits [48], and thus that codon changes occur in the various genome segments. To investigate whether the observed changes in t-loop ΔG stability across the IAV H3N2 genome segments were the result of codon mutation, we plotted the change in ΔG against the amino acid changes (Fig. 5A) for the PB2 (14 changes), PB1 (13 changes) and PA (21 changes) segments (Fig. 5B). Quantification of the number of t-loop ΔG changes that co-occurred with amino acid changes showed that approximately 69% of t-loop ΔG changes occurred independently of changes at the codon level in segment 2 (Table 1). For segment 1 and 3, 86-81% of t-loop ΔG changes occurred independently of changes at the codon level (Table 1). Taking into account the number of codons encoding the three RNA polymerase subunits (i.e., 2232) and the number to t-loop ΔG changes we observe relative to the 1968 H3N2 isolate, a chi-squared analysis yields p = 0. Overall, this suggests that the t-loop ΔG changes occur independently of non-synonymous mutation.

**Figure 5.**
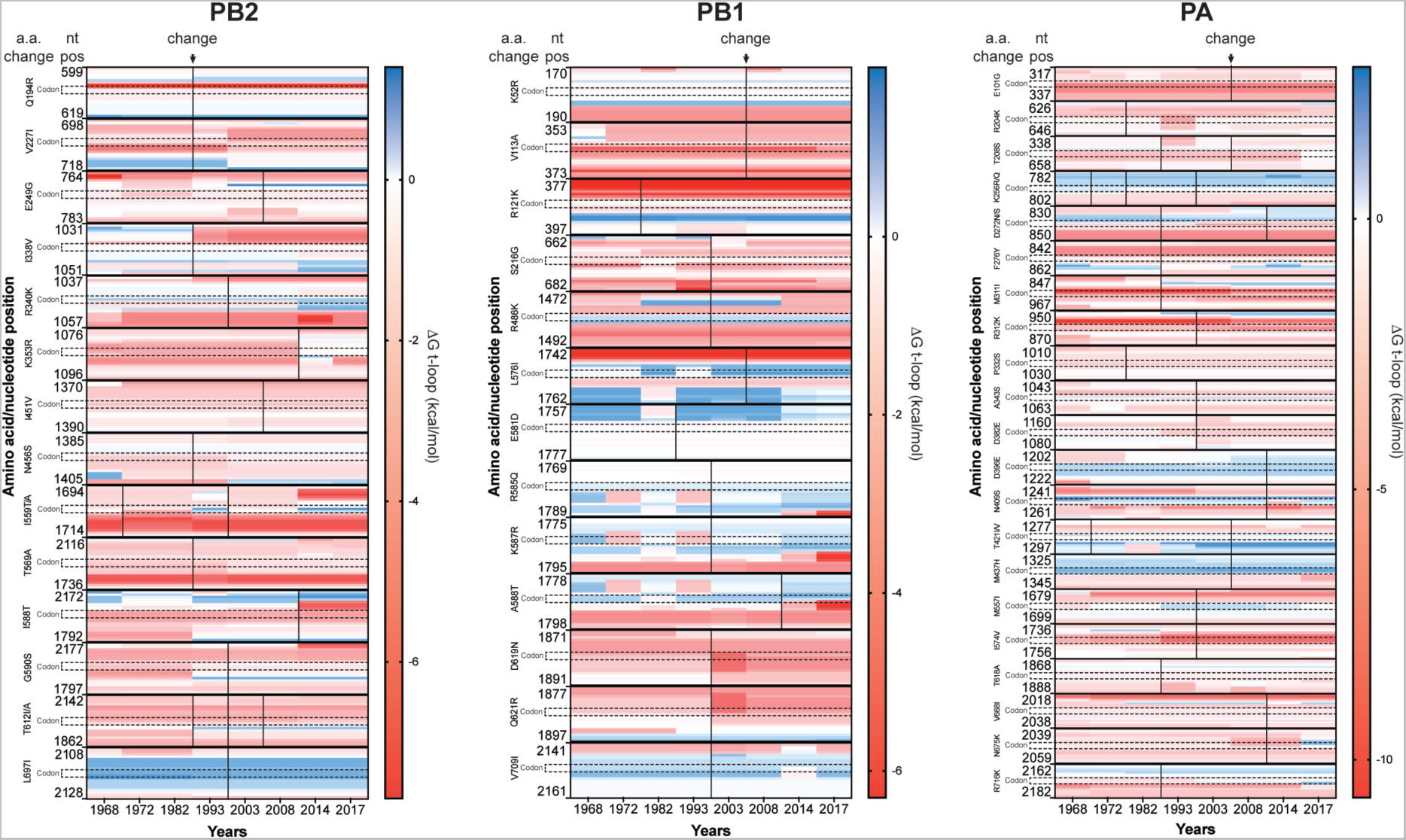
Co-occurrence of codon and t-loop stability change. The IAV RNA polymerase genes accumulate codon changes that alter the protein amino acid sequence of the RNA polymerase subunits. These amino acid changes alter protein function and RNA polymerase activity. To investigate if the observed trends in t-loop destabilization were driven by changes in the codons, we plotted the t-loop stability for the codons (annotated vertically with dotted boundaries) that change for each of the RNA polymerase genes over time and 17 surrounding nucleotides. The color scale shown next to each graph indicates the t-loop ΔG in kcal/mol. Codon changes are indicated with a solid vertical line. A further vertical line is shown as an additional change occurred. Quantification of the co-occurrence of codon and t-loop stability changes is shown in Table 1.

**Table 1:** Coincidence of t-loop stability changes and codon mutation.

| Segment | Codon mutation co-occurrence |  | Total # of codons | Total # of t-loop changes* |
| --- | --- | --- | --- | --- |
|  | Yes # (%) | No # (%) |  |  |
| Segment 1 (PB2) | 2 (14.3 %) | 12 (85.7 %) | 759 | 1697 |
| Segment 2 (PB1) | 4 (30.8 %) | 9 (69.2 %) | 757 | 1649 |
| Segment 3 (PA) | 4 (19 %) | 17 (81 %) | 716 | 1600 |

\*t-loop change is defined as any change in the average $\Delta G$ over the years after 1968 relative to 1968.

### The IAV H1N1 genome shows a loss of secondary structure destabilization

The above data showed that the putative t-loops in the H3N2 PB1 segment were destabilized more quickly than the t-loops in the H3N2 PA and PB2 segments. We wondered if the different origin of the PB1 segment, which had been introduced into IAV H3N2 through a reassortment event with an IAV from the avian reservoir prior to 1968, was driving this increased rate in transient RNA structure destabilization [49]. To explore the possibility that adaptation of an avian origin genome segment to the human host was contributing to the change in t-loop stability in the PB1 segment, we analyzed the genome sequences of IAV H1N1 isolates (Fig. 6).

**Figure 6.**
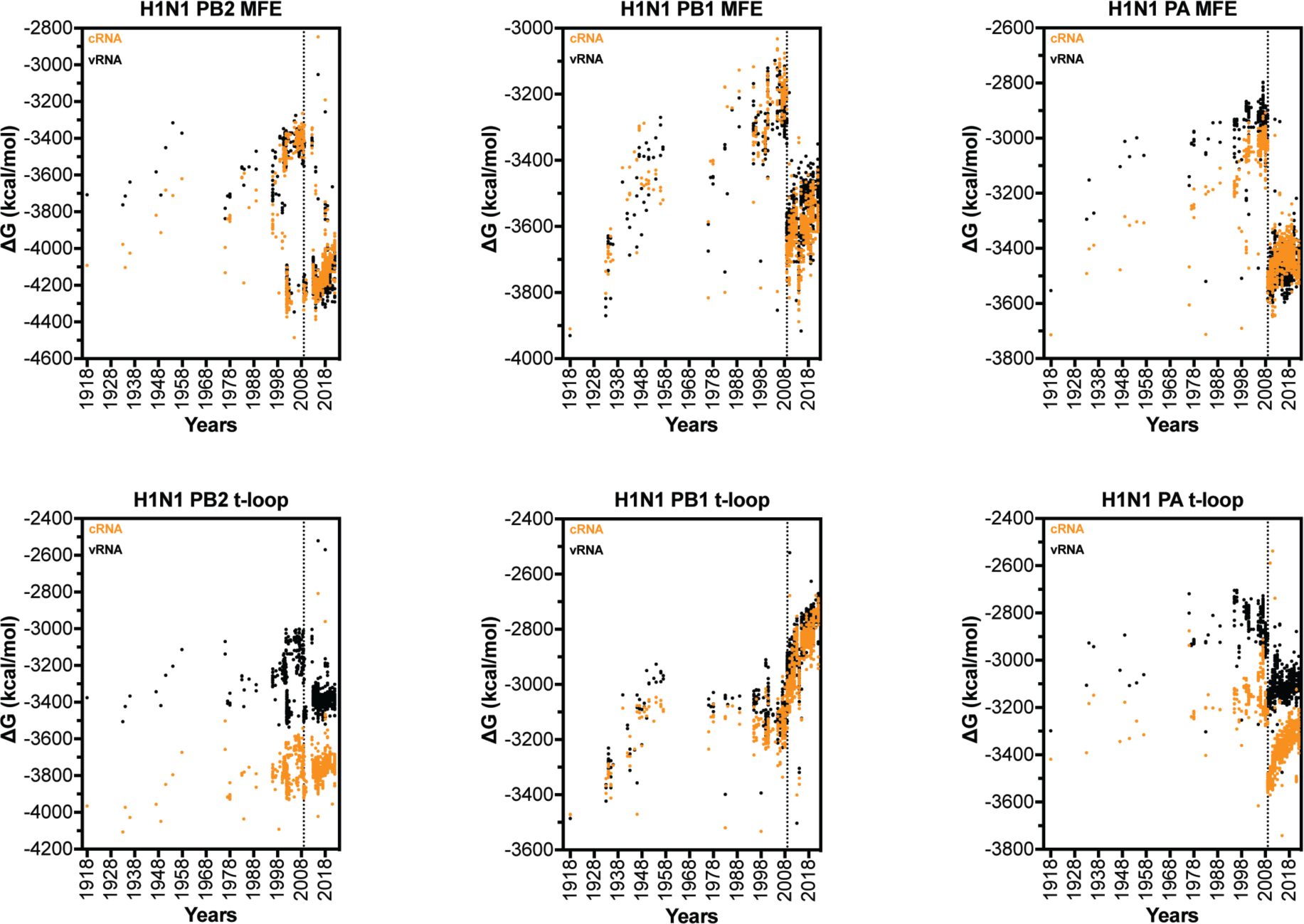
Total MFE and t-loop free energy analysis for IAV H1N1 isolates between 1918 and 2021. Analysis of the cRNA (orange) and vRNA (black) is shown. For each year, the first 50 full-length sequences identified were analyzed and plotted. Dotted vertical line indicates start of 2009 H1N1 pandemic.

In recent memory, H1N1 IAV has been associated with two pandemics in which one or more genome segments were of avian IAV origin [50,51]. In 1918, an avian IAV spilled over into humans and gradually adapted, until it stopped circulating in humans at the start of the 1957 H2N2 pandemic. The same partially adapted H1N1 strain then remerged in 1977 and continued to circulate as seasonal IAV. In 2009, a triple-reassorted pandemic virus emerged, which contained PB2 and PA segments from an avian IAV and a PB1 segment from a human-adapted H3N2 IAV [52,53].

Analysis of the t-loop stability in the three longest H1N1 IAV genome segments revealed a gradual loss of overall t-loop stability in all three segments between 1918 and 1957. Between 1977 and 2009, the t-loop stability remained relatively unchanged for the PB1 segment, whereas the PA segment continued to show some destabilization. The t-loop stability in the PB2 segment was more diverse, and particularly wide-spread from 1998 to 2009. In 2009, a shift in t-loop stability occurred in the PA segment, in line with our expectations for the introduction of an avian-adapted gene segment through reassortment. At the same time, the H3N2-derived PB1 segment displayed a rapid reduction in t-loop stability, whereas the avian-adapted PB2 segment showed only a modest reduction in t-loop stability. Analysis of the overall MFE ΔG in the H1N1 genome segments showed that introduction of a new segment also caused a shift in MFE ΔG. In contrast to the t-loop ΔG, no trend in MFE ΔG change was observed following the reassortment event. In line with our analysis of the H3N2 genome, this suggests that t-loop adaptation occurs independently of changes in other RNA structures in the genome. Overall, the above analysis suggests that IAV reassortment is followed by changes in t-loop stability, and that these changes can occur in both avian-adapted as well as human-adapted IAV genome segments following reassortment.

### The influenza B virus genome does not show secondary structure destabilization

Having observed the relatively rapid change in IAV H3N2 and H1N1 secondary structure stability following IAV reassortment, we analyzed the genome of the influenza B virus (IBV) (Fig. 7). Since current IBV strains have been co-circulating among humans and not been associated with spill over events from a non-mammalian animal reservoir, we expected no (rapid) change in t-loop ΔG over time. Analysis of the t-loop stability indeed showed no trend that was indicative of stabilization or destabilization. The lack of a trend was both evident for both the vRNA as well as the cRNA sense of the three segments investigated (Fig. 7).

**Figure 7.**
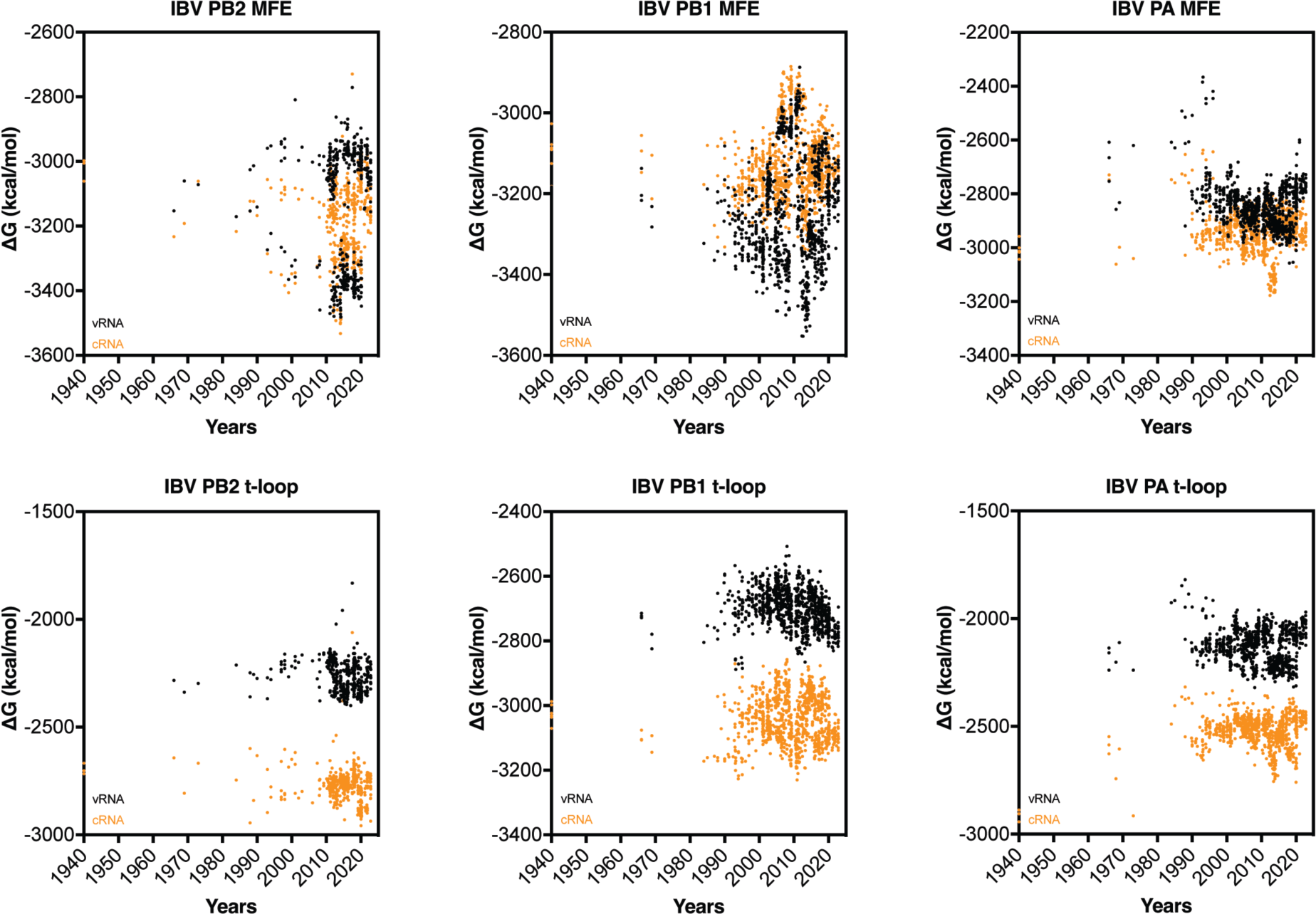
Total MFE and t-loop free energy analysis for IBV. Analysis of the cRNA (orange) and vRNA (black) is shown. For each year, the first 50 full-length sequences were analyzed and plotted.

In contrast to the t-loop analysis, analysis of the MFE showed a more complicated picture for the first three segments of the IBV genome (Fig. 7). For both the PB2 and PB1 segments, we noted a bifurcation in the MFE stability around 1990. This bifurcation, which reduced and increased stability, was particularly prominent for the vRNA sense, and likely linked to the two IBV lineages that have been co-circulating since the 1980s. The MFE of the PA segment was more broadly distributed and did not show a pattern. Overall, these analyses indicate that the t-loop stability in the IBV genome segments is different from the IAV genome segments. It is tempting to speculate that this difference between IAV and IBV is caused by the absence of spill over of IBV strains from a non-mammalian animal reservoir.

### The SARS-CoV-2 genome shows a loss of secondary structure destabilization

In 2019, SARS-CoV-2 emerged as the causative agent of the COVID-19 pandemic. Over the course of the pandemic, point mutations and deletions occurred and accumulated in the SARS-CoV-2 genome. Some of these mutations have been linked to amino acid changes that improved host receptor binding, while others caused deletions of known secondary structures. However, many synonymous mutations and less obvious deletions have not yet been linked to improved viral functions, although it has been observed that some variants display increased replication in tissue culture, suggesting that these mutations and deletions may contribute to viral replication efficiency [54]. Based on our observations with IAV, we hypothesized that t-loop-like structures may be changing in the SARS-CoV-2 genome.

To investigate the above question, we first estimated the footprint of the SARS-CoV-2 replication and transcription complex (RTC). To this end, we aligned motif C of the cryo-EM structure of the SARS-CoV-2 RTC (PDB 7CYQ) with motif C of the cryo-EM structure of the IAV RNA polymerase (PDB 6T0V) in Pymol and estimated the route that template RNA would take through the polymerase. We concluded that 20-22 nt of the template would be covered by the RTC during RNA synthesis [41,55]. Our initial analyses showed that the above footprint parameter range did not affect the trends for the calculated MFE ΔG and the t-loop ΔG. We therefore set the footprint to the lower of these values for the data presented below, in order to collect more t-loop datapoints per genome.

We next analyzed the original SARS-CoV-2 isolate and several SARS-CoV-2 variants of concern using our computational model. For each variant we analysed the first 50, full-length, high-quality sequences published. We found that both in the positive and negative sense of the full-length genome, the MFE ΔG and t-loop ΔG trended towards a less negative ΔG, and thus less stable RNA structure content, over the course of the COVID-19 pandemic (Fig. 8). The trends could be fit with linear regression (p ≤ 0.0001).

**Figure 8.**
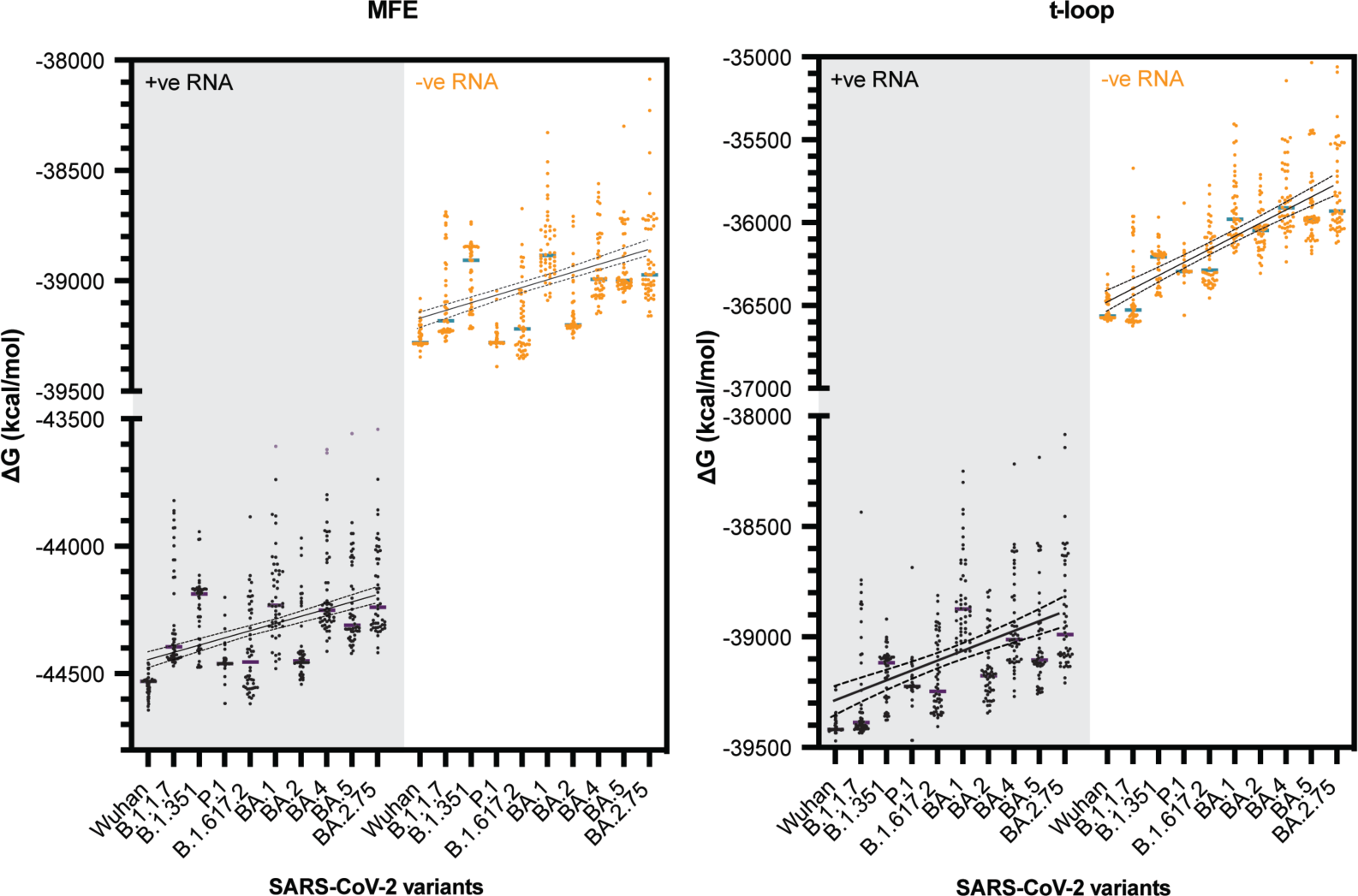
Total MFE and t-loop free energy analysis for SARS-CoV-2. Analysis of the positive sense genomic RNA (black; purple bars indicate mean) and negative sense RNA (orange; green bars indicate mean) is shown. For each year, the first 50 full-length sequences were analyzed and plotted.

Unlike IAV strains, SARS-CoV-2 has only been circulating among humans for a short time. To investigate whether the observed trends were generated by chance, we simulated mutations in the reference SARS-CoV-2 genome, with frequencies of 1.8 x 10^-3^ and 3.95 x 10^-4^ per nucleotide per year as the upper and lower bounds of the whole genome mutation rate [56,57]. We performed the mutation simulations for 3 consecutive years, while maintaining the codon sequence, and next calculated the t-loop 1′G for the *in silico* variants. As shown in Fig. S3, the t-loop ΔG for the *in silico* variants was strikingly less negative than the actual SARS-CoV-2 variants, suggesting that the observed changes in the analyzed SARS-CoV-2 variants had not likely arisen by chance, but also that the changes in the SARS-CoV-2 genome are more constrained than assumed during *in silico* mutation. This result contrasted the *in silico* mutation of the IAV segments.

## Discussion

Respiratory RNA viruses evolve as they circulate among humans. It is well understood how mutations in viral proteins affect enzyme function and protein-protein interactions. In addition, mutations in long-range RNA secondary structures may affect RNA genome packaging or genome circularization during positive sense RNA virus replication. Currently little is still understood about the role of transient RNA secondary structures, or t-loops, which were proposed to only form when the viral replication complex copies the viral genome and modulates the local RNA secondary structure [40].

We here considered the role of the IAV RNA polymerase and NP footprint to compute putative t-loops in the IAV genome segments PB2, PB1 and PA. We observed that the transient RNA structure landscape starts to evolve when emerged IAV strains circulate in humans. Interestingly, we also observed that the RNA polymerase of a 2017 H3N2 IAV isolate was more sensitive to t-loops than the RNA polymerase of a 1968 H3N2 IAV isolate, which suggests that the transient RNA structure landscape and t-loop sensitivity of the RNA polymerase are matched. We observed a similar trend in the evolution of SARS-CoV-2, which recently emerged as a zoonotic virus from an unknown animal reservoir. Such a trend was not evident in the IBV genome, which has circulated in humans without the introduction of non-mammalian adapted sequences for decades. Possible drivers for the observed trends in IAV and SARS-CoV-2 include differences in cellular environment between hosts, including differences in the temperature at the site of replication [58].

A bias in mutations towards bases not involved in base pair formation has been previously observed [59]. It is possible that the change in the transient RNA structure landscape provides a fitness advantage to emerging respiratory RNA viruses, either by increasing the efficiency of viral replication or modulating the volume of aberrant viral products formed during viral RNA synthesis. In turn, the increased efficiency of viral replication could limit the detection of viral RNA by host pathogen receptors, such as RIG-I and MDA5, and contribute less to the severity to clinical disease. In addition, a reduction in RNA secondary structure (more equivalent to the MFE data in our analysis) has been proposed to reduce recruitment of human proteins, especially those found in stress granules [60].

Our observations were almost unilaterally based on *in silico* models that were derived from experiments performed on short IAV templates [40]. In vitro assays will need to be performed for the SARS-CoV-2 RNA polymerase to confirm that this enzyme is indeed sensitive to the structures studied here. In addition, cell-based tools will need to be developed to confirm the existence of t-loops in IAV and SARS-CoV-2 genomes and test our findings. Our *in silico* model also has putative limitations, because it does not account for secondary RNA structures that reside between NP molecules or structures that may be stabilized by RNA-protein interactions in a cellular environment. Such RNA-protein interactions may modulate viral replication processivity in addition to t-loops. Our model also fails to account for the possibility of multiple RNA conformations, instead relying on prediction of the most thermodynamically stable structure. Future work will need to be performed, if possible, to map such potential elements and their variation among viral strains and test hypotheses our study proposes. Future analyses can also focus on developing new statistical frameworks for the rate of stabilization vs destabilization of RNA elements in viral genomes.

Even with the above caveats in mind, our results suggest that changes in transient secondary structure stability are occurring as emerging respiratory RNA virus adapt to humans. This implies that these elements affect viral replication and that a reduction in stability provides a fitness benefit for replication in human hosts. Removing elements from the viral genome that affect processivity and result in early termination or recombination may increase the efficiency of viral replication. Along the same lines, an increase in stability should provide a fitness advantage in non-mammalian hosts, should a human-adapted virus spill back into a non-mammalian host. Together, these results advance our insight into respiratory RNA virus evolution, and we believe that they yield new testable hypotheses that could help strengthen our framework for studying respiratory RNA virus replication and adaptation.

## Material and methods

### Cells and plasmids

Human embryonic kidney 293T (HEK293T) cells were originally sourced from ITCC and routinely checked for mycoplasma infection. HEK293T cells were maintained in DMEM containing 10 % FBS, glutamate, pyruvate, and high glucose (Gibco). WSN pcDNA3-based protein expression plasmids and pPol-based viral template RNA expression plasmids were described previously [61–63]. The open reading frames of the H3N2-2017 and H3N2-1968 PB2, PB1, PA and NP genes were cloned into the pPPI4 expression vector using Gibson cloning. The firefly luciferase reporter plasmid under the control of the *IFNB* promoter [pIFΔ(−116)lucter] and the transfection control plasmids constitutively expressing *Renilla* luciferase (pTK-*Renilla*) were described previously [63].

### Transfections and IFN reporter assays

Transfections to perform RNP reconstitutions and IFN-β reporter assays in HEK293T cells were essentially performed as described previously [63]. RNP reconstitutions were carried out in a 24-well format by transfecting 250 ng of the pcDNA or pPPI4 plasmids encoding PB1, PB2, PA, NP and 250 ng of pPolI plasmid encoding a mvRNA template under the control of the pPolI promoter. HEK293T cells were additionally transfected with 100 ng of plasmid expressing firefly luciferase from the IFN-beta promoter and 10 ng of plasmid expressing *Renilla* luciferase. Transfections were performed with lipofectamine 2000 (Invitrogen) to the manufacturer’s specifications. Twenty-four hours later, the medium was aspirated, cells were washed with 1 ml PBS and split into two fractions. Cells were pelleted at 2,500 rpm using a benchtop centrifuge. The pellets from one fraction were resuspended in 30 μl of 1x Laemmli buffer for western blotting and the other fraction was resuspended in 50 μl PBS for the IFN-β promoter activity assay. The assay was done in duplicate using 25 μl of cell suspension in PBS per well in a white 96-well plate format. Next, 25 μl of DualGlo reagent (Promega) was added per well, samples were incubated at RT for 10 min (in dark), and the firefly luciferase readings were taken using a Synergy LX Multimode Microplate Reader (Biotek). Twenty-five μl of Stop-Glo reagent/well was added next, plate was incubated for 10 min at RT (in dark), and the *Renilla* luciferase readings were taken. Firefly luciferase values were normalized by the *Renilla* luciferase values. Data analysis was performed in Prism 9.5.0 (Graphpad).

### Antibodies and Western blotting

IAV proteins were detected using rabbit polyclonal antibodies anti-PB1 (GTX125923, GeneTex), and anti-NP (GTX125989, GeneTex) diluted 1:2000 in blocking buffer (PBS, 5% bovine serum albumin (RPI), 0.1% Tween-20 (RPI)). Cellular proteins were detected using the rat monoclonal antibody anti-tubulin (MCA77G, Bio-Rad) diluted 1:3000 in blocking buffer. Secondary antibodies IRDye 800 donkey anti-rabbit (926-32213, LI-COR) and IRDye 680 goat anti-rat (926-68076, LI-COR) were used to detect Western signals with a LI-COR Odyssey scanner.

### IAV H3N2 and H1N1, and IBV sequence acquisition and processing

Initially, our analysis was performed on a limited, representative set of H3N2 sequences. To expand this set, 50 full-length IAV H3N2 sequences per year were acquired from EpiFlu (gisaid.org) for 1968 – 2014, and 2017 was downloaded from NCBI (see Table S2). Fifty full-length IAV H1N1 per year were acquired from NCBI for 1918 – 2021. The acquired sequences had >90% full gene segment length and <1% ambiguous nucleotide residues in all gene segments. The 11-12 nt-long conserved IAV promoter sequences were added manually if they had been trimmed and/or were missing in the downloaded sequences. Incomplete sequences that could not be manually updated were removed.

### SARS-CoV-2 variant sequence acquisition and processing

Sequences were acquired from EpiCoV (gisaid.org) are listed in Table S2. Briefly, sequences were required to fit the following criteria: high coverage (<1 % N and <0.05 % unique amino acid mutations), human isolate, complete date record, and considered a complete sequence. Sequences were then sorted by sample collection date and the first 50 complete sequences selected for downstream analysis. This approach allowed us to only include the first 50 sequences of a new variant of concern, limiting putative sequence drift, incorrect variant calling, and recombination between variants. A limitation of our data selection procedure was that it overrepresented the geographic regions in which variants of concern emerged.

After selection, sequences were aligned using EMBL-EBLI Clustal Omega [64]. We noted substantial sequence variation at the 3ʹ and 5ʹ ends of the SARS-CoV-2 genome, which was likely caused by differences in amplicon design or genome assembly among labs. Prior to further analysis, we trimmed the genome sequences to nucleotide positions 602 and 27,552 of the SARS-CoV-2 reference sequence (NC_045512.2). The resulting sequences were approximately 26935 nt in length, with variation stemming from insertions or deletions.

### *In silico* mutation of the SARS-CoV genome

The SARS-CoV-2 reference genome (NC_045512.2) was mutated at frequencies of 1.8 x 10^-3^ and 3.95 x 10^-4^ per nucleotide per year [56,57]. These frequencies were used as upper and lower estimations for SARS-CoV-2s whole genome mutation rate. IAV H3N2 PB2 and PB1 genome segments were mutated at frequencies 3.2 x 10^-3^ and 2.5 × 10^-4^ to represent upper and lower bounds of estimated mutation rates [46,47]. Mutations were performed in python

3.10.0 using the Mutation-Simulator package [65].

### In silico analysis of t-loop and other secondary RNA structures

Our t-loop analysis was essentially performed as described previously [40], but with minor modifications to account for NP binding up and downstream of the IAV RNA polymerase during replication and RNP assembly (Fig. 2). Specifically, we used a sliding window analysis, in which a sequence covered by the 20-nt footprint of the IAV RNA polymerase is blocked from participating in secondary RNA structure formation. Our *in silico* analysis starts with the RNA polymerase binding to the 3’ end of a viral genome (segment), which places residue 16 within the 20-nt footprint in the active site [41,42]. We next assume that NP has a footprint of 24 nt, and that the separation of the template RNA from the downstream or upstream NP in the RNP makes 24 nt available for base pairing (Fig. 2B) [43]. With both an upstream and downstream NP are removed, t-loop as well as any other secondary RNA structure formation can be calculated between the two 24-nt stretches.

The above sliding window was started at the 3’ end of the viral genome and moved in 1-nt intervals along the length of the template RNA. For each window, the RNA sequence upstream of the RNA polymerase footprint, downstream of the RNA polymerase footprint, and between the upstream and downstream sequences was analysed and the stability ΔG of the RNA structures calculated using the ViennaRNA package v.2.5 within Python 3.9 [66]. Locations were recorded based on the position of the active site (i.e., starting at position 16 with ΔG>0, because no template had emerged from the RNA polymerase yet). Next, the total ΔG that the RNA polymerase encounters while copying the template was calculated.

### Consensus sequence generation and GC content analysis

H3N2 consensus sequences for PB2, PB1 and PA were generated using EMBOSS Cons and GC content was calculated using Bio.SeqUtils package from BioPython [64,67]. The python script is provided in the Supplemental Methods.

## Acknowledgments

The authors would like to thank Dr. Michael Oade, Rene Vigeveno, and Sarah van Leeuwen for discussions and reagents. Portions of the work reported in this paper were performed using the Princeton Research Computing resources at Princeton University, which is a consortium of groups led by the Princeton Institute for Computational Science and Engineering (PICSciE) and Office of Information Technology’s Research Computing. CVR was supported by a studentship from Public Health England. KRS was supported by NIH grants R01 GM140032 and R01 AI170520. KB was supported by NIH grant DP2 AI175474. ATV was supported by NIH grants R21 AI147172, DP2 AI175474 and R01 AI170520, and Wellcome Trust and Royal Society grant 206579/17/Z.

## Supplemental information

Figures: 3

Tables: 2

## Notes

### Competing Interest Statement

The authors have declared no competing interest.

### Summary of Updates

Updated Fig. 1, 3-8 for clarity. Added 3 supplemental Figs with additional analyses. Clarified analysis in main text.

