## Supplemental Figures for "Evolution of transient RNA structure-RNA polymerase interactions in respiratory RNA virus genomes"

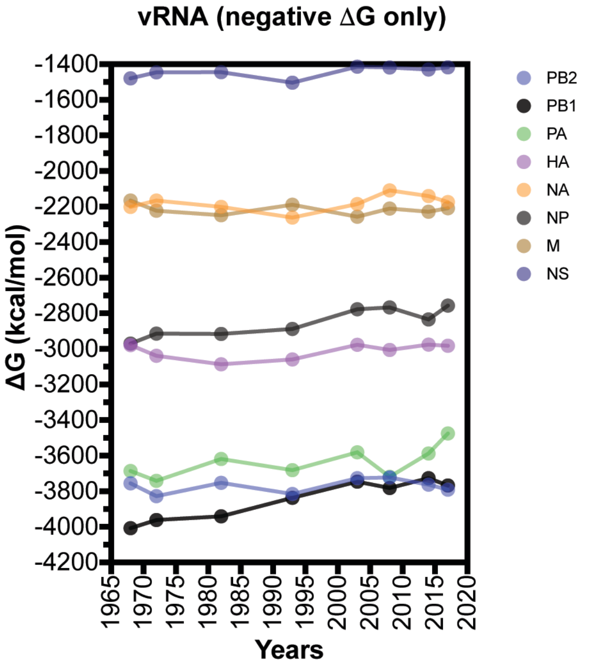


**Figure S1. Analysis of t-loop free energy for IAV H3N2 isolates between 1968 and 2017 using negative free energy values only.**

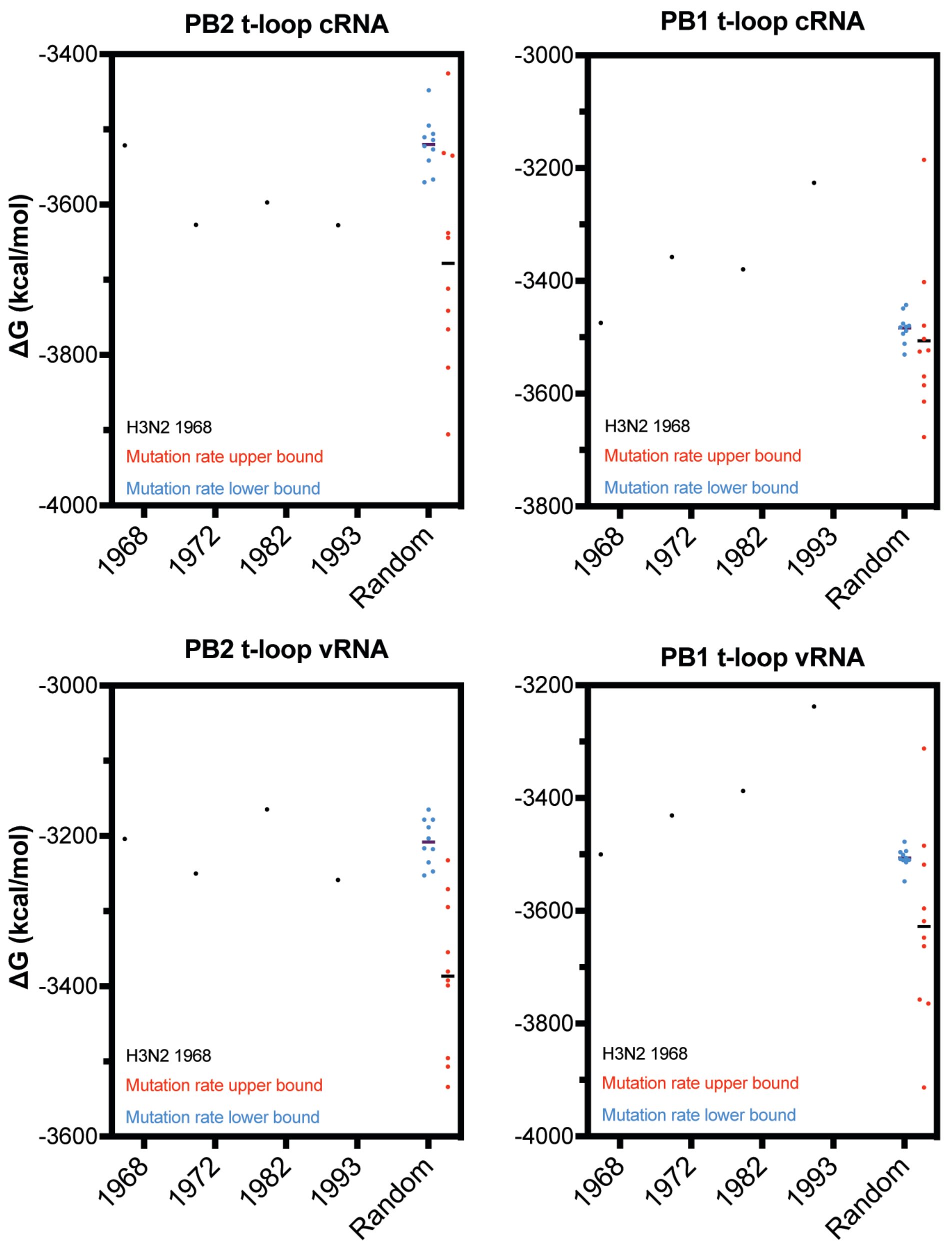


**Figure S2. In silico mutation of IAV genome segments**. Comparison of the t-loop free energy in the IAV PB2 and PB1 encoding genome segments (vRNA sense) following random mutation (in cRNA sense to preserve codon information).

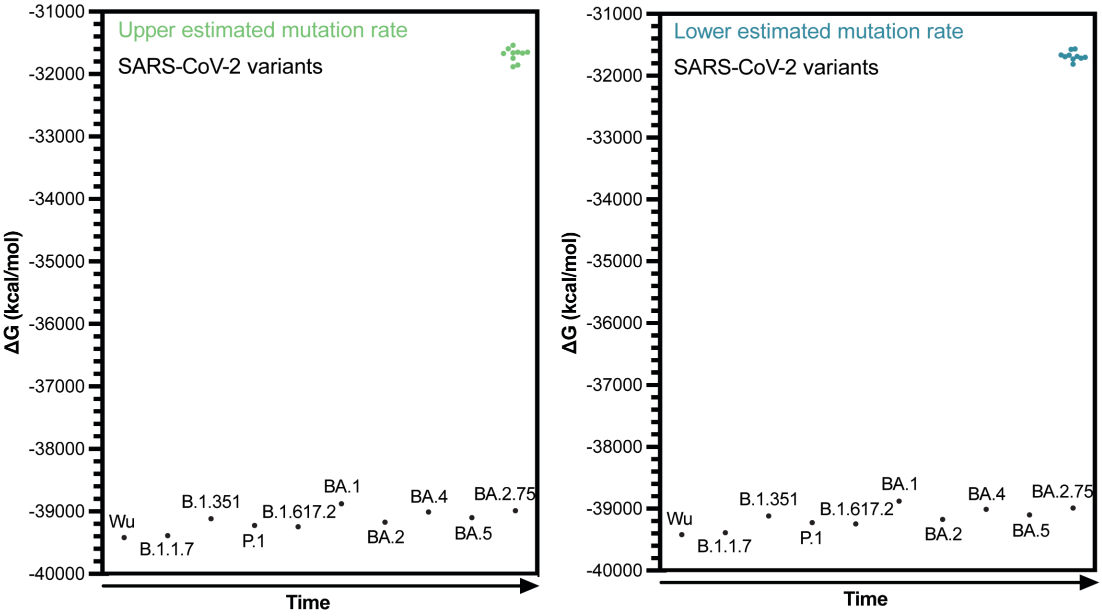


**Figure S3. In silico mutation of SARS-CoV-2 genome**. Comparison of the t-loop free energy in the SARS-CoV-2 genome following random mutation at two different rates of the Wuhan strain relative to the mean observed t-loop free energy in SARS-CoV-2 isolates.
